# Minute- and second-scale hippocampal network dysfunctions in the 3xTgAD mouse model of Alzheimer’s disease are prevented by TSPO knockout

**DOI:** 10.1101/2025.11.11.687784

**Authors:** Benjamin Vidal, Aurélien M. Badina, Violina Dorogan, Gabriel Schirmbeck, Eduardo Sanches, Kelly Ceyzériat, Marit Knoop, Laurene Abjean, Stergios Tsartsalis, Stephane Sizonenko, Olivier Baud, Philippe Millet, Valerio Zerbi, Benjamin B. Tournier

## Abstract

The 18kDa translocator protein (TSPO) is widely recognized as a biomarker of neuroinflammation, but recent evidence suggests that it may also play a direct role in the pathophysiology of neurological conditions with an inflammatory component such as Alzheimer’s disease (AD). In this study, we leveraged functional ultrasound imaging (fUSi) to assess how TSPO knockout influences brain activity and connectivity in the 3xTgAD mouse model of AD. Resting-state scans were performed in 16-month-old male and female mice across four genotypes: WT, TSPO^-/-^, 3xTgAD or 3xTgAD.TSPO^-/-^. Using a data-driven approach, we identified key functional networks, including a bilateral hippocampal-amygdala limbic network. Dual regression analysis revealed increased engagement of the ventral hippocampus within the limbic network and elevated hemodynamic variance in 3xTgAD mice compared to WT. Moreover, co-activation patterns (CAPs) analysis showed abnormal activation/deactivation cycles involving the dorsal hippocampus, both of which were prevented by TSPO knockout without affecting amyloid or tau burden. Our findings highlight a functional role for TSPO in shaping network dynamics and suggest that its deletion can mitigate brain dysfunction in AD models.

## Introduction

Alzheimer’s disease (AD) represents a major challenge for global public health^1^, and current treatments remain mostly focused on alleviation of symptoms rather than the pathological mechanisms, while lacking efficacy and safety^2^. The new generation of disease-modifying treatments targeting the amyloid cascade have yielded mixed results so far^3^, which justifies the exploration of alternative therapeutic strategies, such as modulation of inflammation which is a major component of the disease^4^. Amyloid β (Aβ) and tau protein accumulation are the first neuropathological markers to appear and have been shown to synergistically induce neuroinflammation through reactivity of glial cells^5–7^. In turn, this inflammation further renders the glial cells reactive, recruits peripheral immune cells and overall contributes to neuronal death^8–10^. Reactive astrocytes and microglia are increasingly studied as actors of the pathology, contributing to decreased neuronal metabolic support, synaptic disruption, loss of oligodendrocyte and myelin sheathing^11–14^.

Among molecular targets, the 18 kDa translocator protein (TSPO), widely used in PET/SPECT imaging as a clinical biomarker for neuroinflammation due to its upregulation in reactive glia^15,16^, has recently emerged as a direct effector in the inflammatory cascade. TSPO density has been shown to increase early in the disease, appearing before dementia in humans and even prior to plaque formation in animal models^17–19^. As the pathology progresses, TSPO levels rise further and correlate with amyloid burden^20,21^. While the exact biological function of TSPO is not fully elucidated, recent evidence suggests it is directly involved in astrocytic reactivity^22^, which plays a crucial role in the inflammatory response in the brain during AD^23,24^, and in the appearance of AD markers. Indeed, in the 3xTgAD mouse carrying the key human mutations PS1M146V, APPSwe, and tauP301L^25^, we previously showed that genetic deletion of TSPO reduces astrocyte reactivity^26^. Importantly, TSPO deletion prevented the memory decline induced by tau overexpression, showing an active role in AD progression rather than being a simple marker^26^.

Since TSPO is absent or only minimally expressed by neurons^27^, the impact of its inhibition or overexpression in glial cells on neuronal activity remains unknown, thereby limiting our understanding of the neural circuit dysfunctions driving cognitive impairments. In this context, examining functional network dynamics measured as brain-wide patterns of connectivity and activity offers a sensitive and translational biomarker of neural health, circuit integrity and disease progression^28^. Importantly, brain network activity emerges from the integration of cellular, molecular, and synaptic changes, capturing a holistic view of brain function that unimodal measurements often miss. Within this framework, time-varying network states alterations appear especially insightful, as they correlate with the progression of AD^29^.

Large-scale neuroimaging techniques such as functional MRI enable non-invasive mapping of cerebral hemodynamics as a window into whole-brain activity and connectivity in both cortical and subcortical structures^30^. More recently, functional ultrasound imaging (fUSi) has emerged as a key technology to monitor the changes in cerebral blood volume (CBV) with high spatiotemporal resolution and sensitivity^31–34^, with the potential to provide novel insight into the network-level neural signature of AD and other neurological disorders.

In the present study, we employed fUSi to dissect how TSPO knockout modulates functional connectivity in male and female 3xTgAD mice at 16 months, a disease stage characterized by widespread amyloid and tau accumulation^35^. Our aim is to elucidate the causal relationship between TSPO and network-level activity alterations, thereby evaluating TSPO as a potential therapeutic target in AD.

## Material and Methods

### Animals

A total of 76 C57BL/6 mice (16 months-old, males and females) were used in this study, divided into 4 groups of different genetic background (WT, TSPO^−/−^, 3xTgAD and 3xTgAD.TSPO^−/−^). Experimental approval was obtained from the ethics committee for animal experimentation of the canton of Geneva, Switzerland. Mice were kept under standard light/dark conditions with *ad libitum* access to food and water. Animals were scanned in random order, and data were acquired, preprocessed, and analyzed blind to the experimental group.

### Surgical procedure

Immediately before the imaging session, animals were sedated and anesthetized with intraperitoneal injections of medetomidine (0.6mg/kg), ketamine (70mg/kg) and buprenorphine (0.1mg/kg), and a local head subcutaneous injection of lidocaine (7mg/kg). The top of the skull was removed using a circular saw for micro-drill (FST #18000-20), and ultrasound transmission gel (sterile Aquasonic 100) was applied before imaging. As the time from the first injection to the end of the imaging session did not exceed 30 minutes, no additional injection of ketamine was performed.

### Functional ultrasound imaging procedure

After the cranial window was performed, mice were placed into a stereotaxic frame for the imaging procedure. Body temperature was continuously monitored during the acquisition. The probe switched between 3 coronal planes at -3.5 mm, -2.5 mm and -1.5 mm from bregma during 10 minutes of resting-state, at a sampling rate of 2.5 Hz and final repetition time of 1.8s between two successive images in the same plane (Fig. 1A).

**Figure 1:**
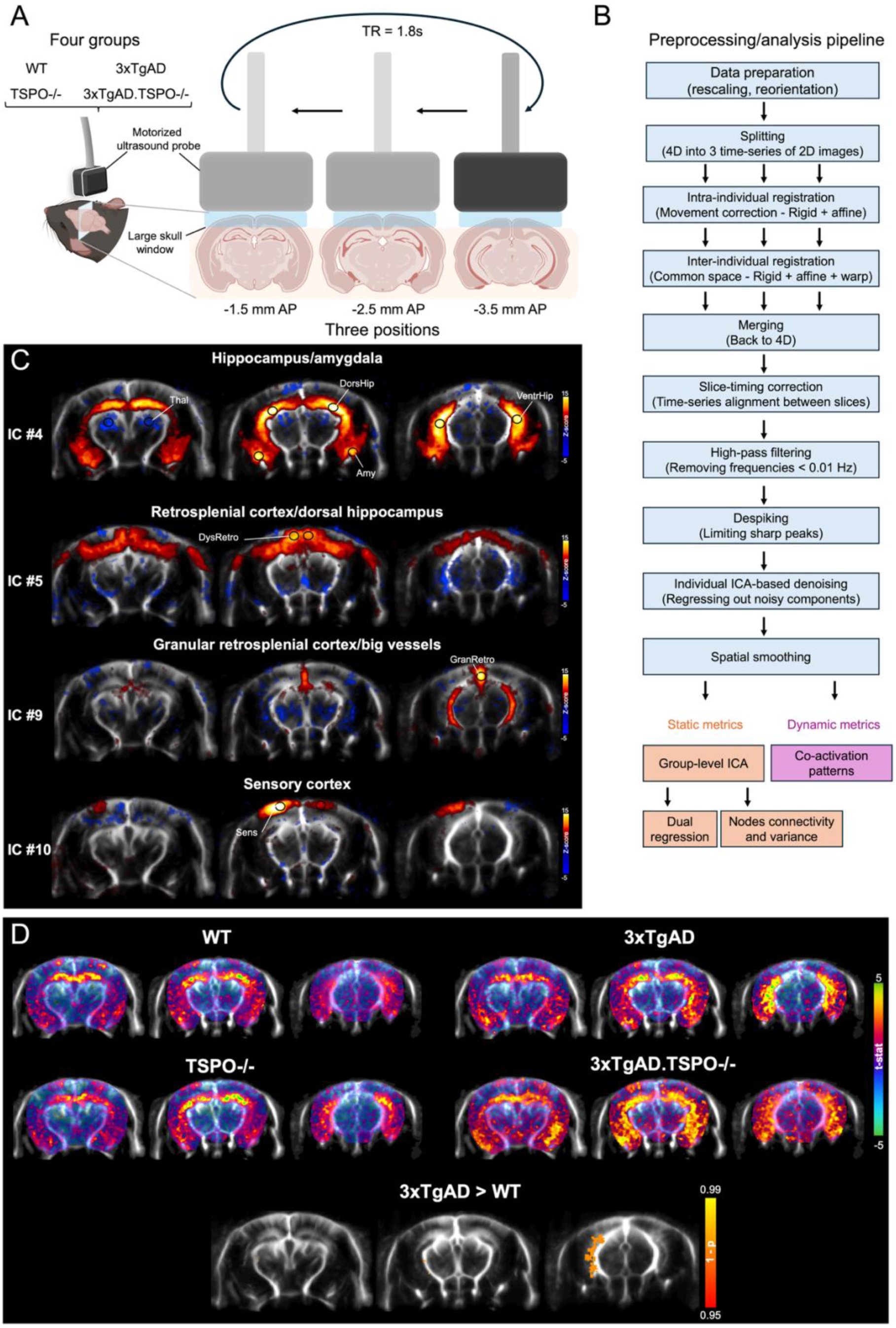
fUSi to probe the effects of TSPO knockout on cerebral functional networks in Alzheimer’s disease mouse model. (A) Schematic of the acquisition protocol targeting hippocampal regions over 3 coronal slices, for a final repetition time of 1.8s between two successive images of the same plane. (B) Description of the main preprocessing and data-driven analysis pipeline of the fUSi dataset. (C) Group-level ICA analysis identified 4 networks of interest in the dataset, from top to bottom: a hippocampal/amygdala network encompassing the entire bilateral hippocampus along the dorso-ventral and rostro-caudal axes; a dysgranular retrosplenial cortex/dorsal hippocampal network; a network of major blood vessels, located mostly in the caudal plane inside the granular retrosplenial area and along the ventral hippocampus; a sensory cortex network, located mostly in the middle plane. (D) Top: Group-level spatial maps of IC#4 (hippocampal-amygdala network), after dual regression and permutation testing. Bottom: Between-groups comparison showed a significant increase in 3xTgAD compared to WT located in the left ventral hippocampus (p<0.05 after permutation and cluster-mass FDR correction). Other between-groups comparisons were non-significant.

Doppler vascular images were obtained using a preclinical ultrasound imaging system (Iconeus V1, Paris, France), based on the Ultrafast Compound Doppler Imaging technique^36^. Planes of interest were located using the automated brain positioning system implemented in the imaging software^37^. The coherent summation of backscattered echoes obtained after successive tilted plane waves emissions allowed the reconstruction of a Compound Plane Wave frame^38^. To extract the blood volume signal from the tissue signal, each stack of 200 compounded frames (acquired at 500 Hz frame rate) was processed with a dedicated spatiotemporal filter using Singular Value Decomposition^39^. The final Power Doppler images are directly proportional to the cerebral blood volume, with spatial and temporal resolutions of 100 µm and 0.4s^40^.

### Ex vivo measurements

At the end of the imaging procedure, mice were euthanized using an intracardiac saline perfusion and their brains were removed. The hippocampus was isolated and was frozen in liquid nitrogen and then stored at -80°C until use. Samples were sonicated in a solution of triton (50mM Tris HCl, 150mM NaCl, 1%Triton x100, protease and phosphatase inhibitors 1x, pH=7.4). Following centrifugation (20 000 g, 20 min, 4°C), the triton (Tx)-soluble fraction (supernatant) was collected. The pellet was dissolved in a guanidine solution (5M guanidine, 50mM Tris HCl, protease and phosphatase inhibitors 1x, pH=8; Gu-soluble fraction). A protein assay was performed using the BCA kit to quantify the total amount of proteins. The determination of Aβ40, Aβ42 and tau phosphorylated at T231 was carried out by ELISA using commercial kits (Aβ40 human ELISA kit; Aβ42 human ultrasensitive ELISA Kit and tau [pT231] human ELISA Kit, Thermo Fisher).

### Functional ultrasound imaging data preprocessing

After initial quality control, 14 mice that displayed low cortical signal due to local damage during the surgery were excluded from the analysis, resulting in the inclusion of 62 mice for preprocessing. Multiple steps were followed to enable voxel-wise analyzes of functional connectivity in the different groups, as shown in Figure 1B. Data were first exported into NIfTI format using the Icostudio software (Iconeus, Paris) and voxel dimensions were multiplied by ten using Matlab for compatibility with analysis tools. Images were reoriented to follow the RAI (Right to left/Anterior to posterior/Inferior to superior) orientation using FSL (‘fslswapdim’, ‘fslorient’) and AFNI (‘3drefit’) functions. The 4D volumes were splitted into three 3D stacks of 2D images (one per coronal plane) using ‘fslsplit’, before proceeding to the intra-individual registration, to avoid any possible sagittal tilt introduced during this step. For a few subjects, coronal planes that appeared too caudal or too rostral from the targeted field-of-view were then excluded. The intra-individual registration, enabling to correct for slow positional drift or progressive changes in the shape of the brain that occasionally occurred, was performed using ANTs (Advanced Normalization Tools, ‘antsMotionCorr’ function) using rigid, then affine registration on the mean image computed across time dimension for each subject. For each coronal plane, all subjects were registered into the same space using ANTs (template creation and transformation matrix estimation: ‘antsMultivariateTemplateConstruction2’ function, SyN method, mutual information metric; transformation of the time-series: ‘ants.apply_transforms’, Python). 6 mice were excluded at this step due to failed normalization, resulting in the final inclusion of 56 mice (WT: n=12, TSPO^−/−^: n=17, 3xTgAD: n=16, and 3xTgAD.TSPO^−/−^: n=11, with overall no significant difference in sex distribution across groups: χ²(3,56) = 4.69, p = 0.196). The 3D data were then merged back to 4D using ‘fslmerge’, before slice timing correction using ‘slicetimer’ FSL function to time-lock first and last coronal planes to the middle plane and approximate simultaneous sampling across slices. Data were then high-pass filtered using FSL-FEAT to remove frequencies below 0.01 Hz, and despiked using ‘3dDespike’ from AFNI to limit the amplitude of possible sharp peaks. To further improve dataset quality, an independent component analysis (ICA) was performed on each individual acquisition using FSL-MELODIC (number of components set to 15). The components that were due to electronic artifacts (typically as horizontal or vertical band patterns) or occurring outside of the brain were manually labeled as noise and regressed out from the data using ‘fsl_regfilt’ function (see Supplementary Figure 1A for examples of individual components from the same acquisition; as a precaution, only obvious sources of noise were removed, not ambiguous components). Finally, a Gaussian smoothing (kernel size 0.5) was applied to the images using ‘fslmaths’.

### Functional ultrasound imaging data analysis

A data-driven approach was chosen for analyzing functional connectivity in the different groups of mice using static or dynamic metrics (Fig. 1B).

For assessing static functional connectivity along the 10 minutes of resting-state, a group-level ICA was performed with FSL-MELODIC, using temporal concatenation of all acquisitions from all groups, with background threshold masking and MIGP (MELODIC’s Incremental Group-PCA, a memory-efficient approximation of the full PCA) options disabled. After voxel-wise demeaning and normalization of the voxel-wise variance, data were whitened and projected into an 18-dimensional subspace using principal component analysis. The whitened observations were decomposed into sets of vectors which describe signal variation across the temporal domain (time-courses), the session/subject domain and across the spatial domain (maps) by optimizing for non-Gaussian spatial source distributions using a fixed-point iteration technique^41^. Estimated Component maps were divided by the standard deviation of the residual noise and thresholded by fitting a mixture model to the histogram of intensity values^42^. This enabled to obtain 18 individual components to describe global functional connectivity patterns from the entire mouse population. Components were excluded from further analysis if they (i) did not exhibit bilateral spatial patterns consistent with known functional neuroanatomy (such as unilateral patterns, vertical stripes, or patterns occurring outside of the brain) and (ii) were dominated by global, slice-wise oscillations. The latter consisted in high-amplitude oscillations that were non-stationary in the frequency domain and that tended to synchronize and desynchronize between slices (see Supplementary Fig. 1A for example in single-session ICA, or ICs #1 to 3 of group-ICA in Supplementary Fig. 1B). These peculiar characteristics led us to discard them from further analyses as they are difficult to interpret and may result from non-neuronal phenomena, possibly related to the anesthesia regimen. From these, four components of interest were selected for a dual regression analysis using the ‘dual_regression’ function of FSL and compared between groups using a general linear model and permutation with the ‘randomise’ function (10’000 permutations, cluster mass-based thresholding for multiple comparisons correction at p<0.05, sex factor as a covariate of non-interest). For further exploration, nodes of interest of equal size (14 voxels) were drawn on the spatial maps of the four selected group-level components using ITK-Snap software, for extraction of the time-series using ‘fslmeants’. The time-series were analyzed in terms of signal variance after centering and normalization in percentage of changes relative to the mean signal, and between-nodes connectivity was estimated using FSLNets toolbox in Matlab (full correlation option, between-groups comparison by permutation testing with ‘nets_glm’, 10’000 permutations and a false discovery rate of 5%, sex factor as a covariate of non-interest).

For assessing dynamic functional connectivity at the single-frame level, co-activation patterns (CAPs) were estimated with the TbCAPs toolbox^43^ in Matlab. All acquisitions from all groups and all frames were included before clustering into ten CAPs. Four CAPs of interest were selected for further between-groups comparisons based on the number of episodes per CAP and the transition probability between those CAPs.

### Statistics

For between-groups dual regression and between-nodes connectivity comparisons, non-parametric permutation testing was used as implemented in FSL, with correction for multiple comparisons (at the cluster level for the voxel-wise dual regression). All others analyzes were conducted with Graphpad Prism, using two-way ANOVAs and appropriate post-hoc tests when data followed normal distribution (as assessed by Kolmogorov-Smirnov test, p>0.05), or alternative non-parametric tests recommended by the software (Mann-Whitney tests to compare a single variable between two groups, or multiple Mann-Whitney tests followed by false discovery rate correction of 5% to compare two groups across several conditions).

## Results

### Identification of bilateral networks of interest across all mice

We first aimed to identify common functional connectivity patterns across all groups of animals in a data-driven manner. To this end, we performed a group-level ICA across the entire dataset, which successfully revealed several anatomically meaningful networks (Fig. 1C). Notably, we focused our analyses on four key networks: (i) a bilateral hippocampus–amygdala limbic network (IC#4 – 6.44% of explained variance), extending along the full rostro-caudal axis of the hippocampus and showing anticorrelated activity with the laterodorsal thalamus. (ii) a bilateral network including the retrosplenial cortex and the dorsal hippocampus, displaying negative correlation with the ventral hippocampus (IC#5 – 6.36% of explained variance); (iii) a network at the margin of the ventral hippocampus and in the granular retrosplenial cortex (IC#9; 5.06% variance explained). While this latter component may primarily reflect contributions from large vessels rather than neuronal activity, we retained it as a component of interest given that vascular pulsatility may represent a key pathophysiological feature of Alzheimer’s disease^44^. Finally, we identified (iv) a sensory cortical network primarily encompassing parietal regions (IC#10 – 4.98% of explained variance). Other independent components were not selected since they did not meet the criterion of bilateral co-activation^45^ (such as ICs #6, #7 and #16), or because their spatial distribution did not match anatomically meaningful brain regions, showing either whole-slice effects (ICs #1-3), or patterns resembling imaging artifacts (ICs #8, #11-15, #17 and #18) (Supplementary Fig. 1A-B).

### Ventral hippocampus over-engagement in the limbic network of 3xTgAD mice is normalized by TSPO deletion

Next, we sought to compare the strength of the networks of interest between groups. Whereas networks involving cortical areas and/or big vessels showed comparable strength across all groups (ICs #5, #9 and #10 - p>0.05, FDR corrected,), we found differences in the hippocampal-amygdala network. Specifically, the first-level spatial t-maps suggested differences between 3xTgAD mice and all other groups, with higher t-values localized bilaterally in the caudal ventral hippocampus in this group (Fig. 1D, top). In contrast, 3xTgAD.TSPO^-/-^ mice showed lower t-values in this region, although their spatial maps appeared more widespread than those of the other groups. Non-AD TSPO^-/-^ mice displayed a network pattern comparable to WT mice.

Between-group analysis confirmed a significant increase in network strength specifically in 3xTgAD versus WT controls, localized to the left ventral hippocampus (p = 0.0264, FDR-corrected; Fig. 1D, bottom). All other group comparisons, including 3xTgAD.TSPO^-/-^ versus WT, did not reach significance. Thus, limbic network hyper-connectivity occurred only in the Alzheimer’s disease mouse model expressing TSPO, whereas TSPO deletion prevented this abnormal functional phenotype.

### TSPO-related limbic network abnormalities reflect a focal increase in ventral hippocampal hemodynamic fluctuations

Having identified a focal connectivity alteration in 3xTgAD mice with cluster-mass thresholding, we investigated in detail the dynamic changes of vascular signals across multiple nodes of the limbic network and other networks. To do that, time courses were extracted from anatomically defined nodes on the group-level ICA maps (Fig. 2A). First, we measured connectivity matrices as Pearson’s correlation over the full 10-minute scans (Fig. 2B). While Z-scores tended to be higher between hippocampal and amygdala subregions in 3xTgAD mice, no significant inter-nodal connectivity differences were detected across groups (all p > 0.05, FDR-corrected), confirming that the observed alterations occur mostly at the focal rather than network-wide level. By contrast, signal variance in ventral hippocampal nodes was significantly higher in 3xTgAD mice relative to both WT and 3xTgAD.TSPO^-/-^ groups (Fig. 2C; post hoc Fisher’s LSD: p = 0.0172 vs. WT, p = 0.0195 vs. 3xTgAD.TSPO^-/-^; two-way ANOVA: gene, F(1,50) = 2.916, p = 0.0939; disease, F(1,50) = 3.172, p = 0.081; interaction, F(1,50) = 3.061, p = 0.0863). No differences emerged in dorsal hippocampal nodes (all p > 0.05). Together, these results further indicates that abnormal CBV fluctuations were restricted to the ventral hippocampus in AD mice expressing TSPO.

**Figure 2:**
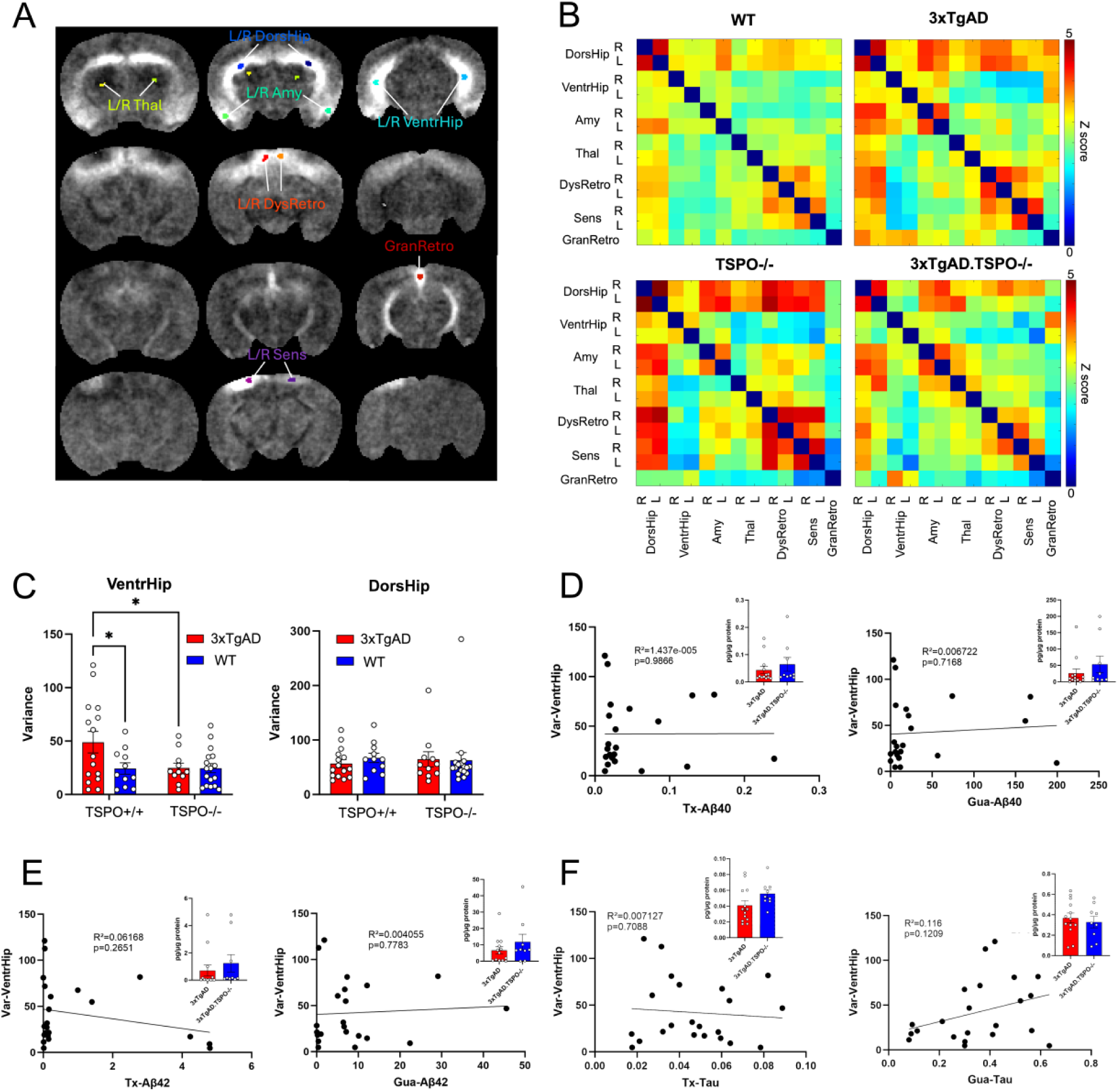
The minute-scale functional alteration is located inside the ventral hippocampus of 3xTgAD mice and prevented by TSPO knockout, independently of amyloid or tau burden. (A) Nodes of interest as defined on the group-level ICA maps (same maps as in Fig. 1C with different colorscale). (B) Mean functional connectivity matrices between nodes of interest defined on the group ICA networks, for each experimental group, expressed as Z-scores. The permutation tests showed no significant differences between groups (p>0.05 after FDR correction). (C) Bar plots (mean +/- SEM) and individual values of the variance of the fUSi signal inside the ventral and dorsal hippocampus show a significant increase in the ventral area in 3xTgAD mice compared to WT and to 3xTgAD.TSPO^-/-^ (Fisher’s LSD test after two-way ANOVA, *p<0.05). (D) Scatter plots of the ventral variance against the concentrations of the soluble and insoluble forms of Aβ40 peptide in the hippocampus, showing no significant correlations. Insets are the bar plots (mean +/- SEM) and individual values of peptide concentrations, showing no significant difference between groups (p>0.05, Mann-Whitney tests). (E) Same scatter and bar plots with the Aβ42 peptide in the hippocampus, also showing no significant correlation with ventral variance and no significant difference between groups (p>0.05, Mann-Whitney tests). (F) Same plots and insets with phosphorylated tau 231, showing no significant correlation nor difference between groups (p>0.05, Mann-Whitney tests).

### TSPO knockout normalizes focal hemodynamic fluctuations without reducing amyloid and tau burden

We hypothesized that the prevention of the abnormal functional phenotype in 3xTgAD mice following TSPO knockout might be attributable to beneficial effects on core pathological hallmarks of AD, namely amyloid and tau accumulation. Given that increased hemodynamic fluctuations in the ventral hippocampus emerged as a signature feature of 3xTgAD mice, we examined whether ventral hippocampal signal variance correlated with levels of poorly and highly aggregated forms of Aβ40 and Aβ42 in the hippocampus of 3xTgAD and 3xTgAD.TSPO^-/-^ animals. No significant correlations were observed (Fig. 2D,E; all p > 0.05, R² < 0.1), and amyloid peptide levels were not lower in 3xTgAD.TSPO^-/-^ than in 3xTgAD mice (Mann–Whitney tests, all p > 0.05). Thus, normalization of the functional phenotype following TSPO knockout could not be explained by a reduction in amyloid burden. We also quantified the hippocampal levels of phosphorylated tau and found no correlation between ventral hemodynamic variance and the soluble or insoluble fractions of p-tau231 (Fig. 2F; all p > 0.05). The soluble form of tau tended to be slightly increased in 3xTgAD.TSPO^-/-^ mice compared to 3xTgAD, whereas the reverse trend was found for the insoluble tau, but all differences were not significant (Fig. 2F, Mann-Whitney test, all p > 0.05). This further indicates that the effects of TSPO knockout revealed by fUSi are not attributable to an overall reduction of the pathological protein burden at this advanced disease stage.

### 3xTgAD mice also display dynamic, TSPO-dependent abnormalities in dorsal hippocampal coactivation cycles

Static connectivity measurements averaged over several minutes can overlook critical features of brain activity, as functional connectivity is now recognized to fluctuate dynamically across rapid transitions between distinct brain states^46^. To capture these dynamics in our mouse models, we performed co-activation pattern (CAP) analysis on all frames across all groups. This approach identified 10 reproducible CAPs (Supplementary Fig. 2), of which 4 were considered relevant for further groups comparisons (Fig. 3A). CAP 1 was characterized by signal decreases in the dorsal hippocampus, retrosplenial cortex, and dorsal thalamus, with concurrent increases in temporal and ventral regions. CAP 2 showed widespread hippocampal and partial amygdala activation, closely resembling the limbic IC#4, along with concurrent cortical suppression. CAP 3 consisted of global cortical activation coupled with subcortical inhibition, while CAP 4 was strongly anti-correlated to CAP 1. The remaining six CAPs corresponded to single-slice global activations (Supplementary Fig. 2), consistent with large-scale waves also observed in the first ICA components. These states reflected three global modes expressed in either positive or negative polarity (“Anti” labels), confirming synchrony–asynchrony effects between slices (Supplementary Fig. 2).

**Figure 3:**
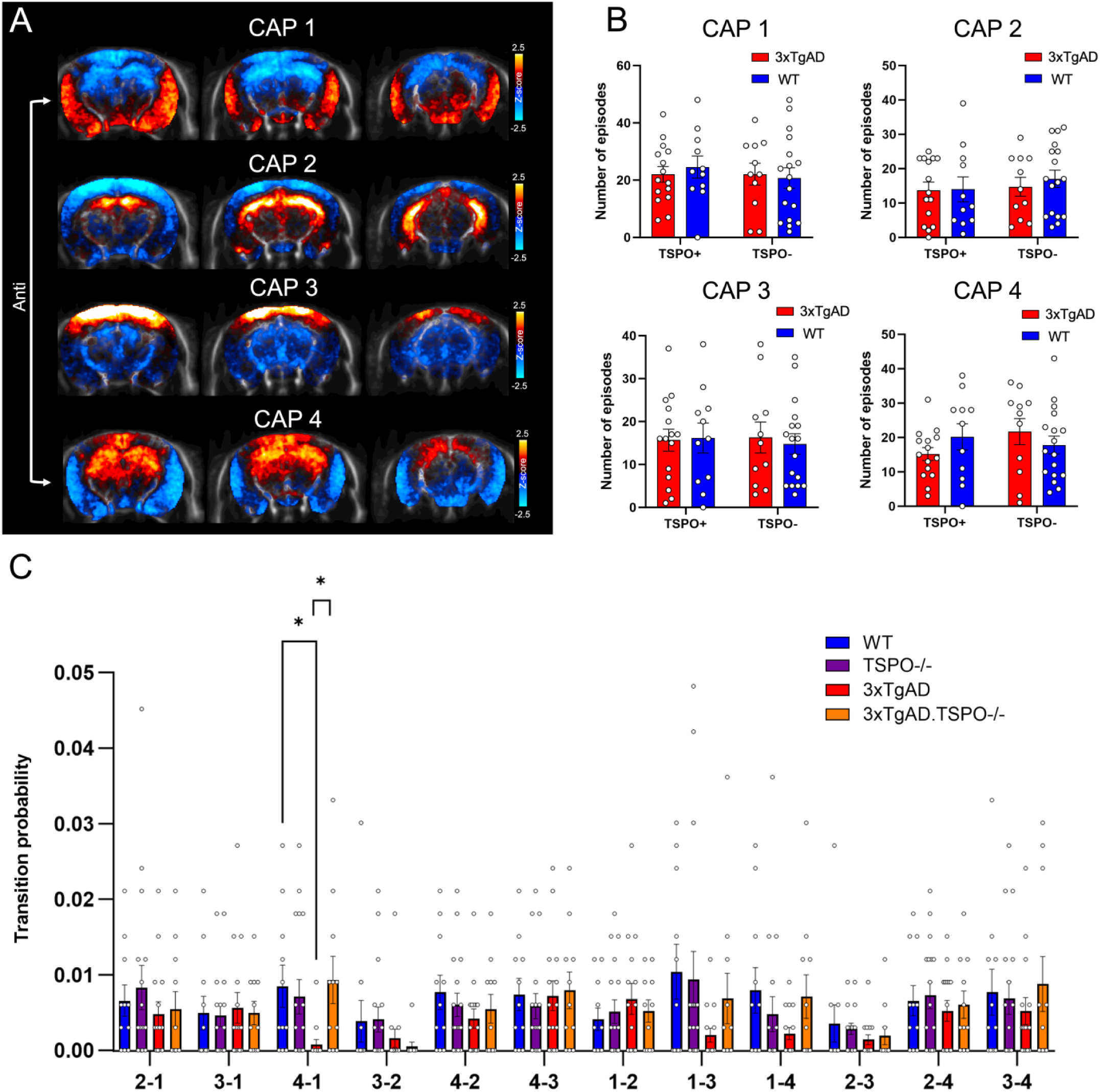
The dorsal hippocampus of 3xTgAD mice displays dynamic alterations of co-activation cycles that are prevented by TSPO knockout. (A) Instantaneous brain configurations of interest, as identified by co-activation patterns analysis performed on all mice and all frames. (B) Bar plots (mean +/- SEM) and individual values of the variance of the number of episodes for each CAP of interest, showing no difference between groups (Fisher’s LSD tests following two-way ANOVA, p>0.05). (C) Bar plots (mean +/- SEM) and individual values of the transition probabilities from one CAP of interest to the others, showing a significant difference in 3xTgAD mice compared to WT and to 3xTgAD.TSPO^-/-^ (*p<0.05, multiple Mann-Whitney tests and FDR correction).

Quantification of CAP occurrences revealed no significant differences across groups (all p > 0.05), indicating that the overall repertoire of brain state dynamics was preserved in AD mice (Fig. 3B). In contrast, analysis of transition probabilities (Fig. 3C) revealed a specific deficit in 3xTgAD mice: the probability of transitioning from CAP 4 to its complementary CAP 1 was significantly reduced compared to both WT and 3xTgAD.TSPO^-/-^ groups (Mann–Whitney tests with FDR correction: 3xTgAD vs. WT, p = 0.0017, q = 0.020; 3xTgAD vs. 3xTgAD.TSPO^-/-^ , p = 0.0018, q = 0.021; all other q > 0.5).

Other group comparisons were not significant (q > 0.87). This indicates that retrosplenial and dorsal hippocampal activation (CAP 4) was less likely to be followed by its reciprocal deactivation (CAP 1) in 3xTgAD mice, suggesting a disruption of this co-activation cycle. This alteration was normalized by TSPO knockout.

## Discussion

Hippocampal dysfunction is a defining feature of Alzheimer’s disease, reflecting specific network-level vulnerabilities that underlie memory decline and cognitive impairment. Yet, how these alterations unfold dynamically across different timescales of brain activity remains poorly understood. Here, by applying functional ultrasound imaging (fUSi) to a well-characterized AD mouse model, we reveal distinctive hippocampal network abnormalities that manifest consistently across both static and dynamic connectivity measures. Importantly, genetic deletion of TSPO prevented these aberrant functional phenotypes, while sparing brain activity in non-AD mice, highlighting TSPO as a critical mediator of Alzheimer’s-related network dysfunction.

Interestingly, our approach identified a distinct alteration along the ventral-dorsal axis in the hippocampal/limbic network. Indeed, the ventral hippocampus displayed a stronger synchronization within the limbic network coupled with an increase in CBV variance over a 10-minute timescale, while the dorsal hippocampus was found to have an aberrant cycle transition at the single-frame timescale. This ventral/dorsal dichotomy is consistent with previous observations in this mouse model. These two regions differ in terms of proteomic and electrophysiological changes, behavioral correlates^47,48^, and vulnerability to pathological protein accumulation during disease progression^26,35^. Major differences have also been reported in the human along the equivalent anterior-posterior axis^49,50^. In 3xTgAD mice, the presence of a dorsoventral gradient in amyloid plaque accumulation could contribute to this functional dichotomy^18^. Conversely, region-specific differences in hemodynamic fluctuations might also influence the dorsoventral pattern of amyloid burden, given their established role in brain clearance^51,52^. Importantly, while CBV fluctuations generally follow neuronal activity^53^, they are not a direct measure of it. The increased hemodynamic variance we observed is paradoxical at this advanced stage of the disease, when reduced neuronal excitability in the hippocampus might be expected. However, this result does not necessarily indicate local neuronal hyperactivity and could be driven instead by glial or vascular mechanisms. Indeed, previous fUSi studies have demonstrated that glial reactivity can lead to a dramatic increase in spontaneous oscillations of CBV during acute neuroinflammation^54^.

In line with this interpretation, we have previously shown that chemogenetic stimulation of hippocampal astrocytes modulates cerebral blood flow in wild-type animals but not in the TgF344-AD rat model^55^, further supporting the growing evidence that astrocyte-vascular coupling is disrupted in AD^56–59^. TSPO knockout has also been reported to reduce astrocyte reactivity in the hippocampus of 3xTgAD mice, thereby restoring normal functional phenotype by decreasing the release of pro-inflammatory cytokines^26^, which may ultimately limit neurotoxicity and prevent aberrant vascular oscillations. Consistently, the same study showed that TSPO knockout restores normal glucose metabolism in 16-month-old 3xTgAD mice^26^. The beneficial effect of TSPO knockout on brain networks and hemodynamics cannot be attributed to a direct effect on amyloid or tau pathology, as neither hippocampal Aβ nor tau burden was reduced. More generally, inter-individual differences in hippocampal hemodynamic fluctuations in the presence or absence of TSPO among 3xTgAD mice were not explained by differences in pathological proteins load. This dichotomy between Aβ/tau burden and restoration of brain activity aligns with previous findings from our lab, where low-dose radiotherapy decreased TSPO levels and inflammatory response and improved memory in TgF344-AD rats without decreasing Aβ load^60–62^, highlighting that inflammation is a key factor in AD symptomatology, beyond these pathological hallmarks^4^.

The specific hippocampal signature bring additional information to the existing literature assessing resting-state functional connectivity in AD models based on hemodynamic changes^63–70,71^. While it is difficult to compare our own results with previous findings due to differences in strains, disease progression, sex, imaging modalities and anesthesia regimen, they are in agreement with previously reported hippocampal alterations^63,67,68,70,71^ and confirm that functional connectivity is impaired even on the shortest timescales of a few seconds^72,73^, emphasizing the importance of dynamic functional connectivity assessment in neuroimaging studies^46^. Here, a single time point was chosen because of the focus on TSPO, but future studies should aim for longitudinal monitoring of disease progression (as previously done in fMRI for the double transgenic App^NL-G-F^/hMapt model^70^) using chronic cranial windows^74,75^, with the possibility of avoiding anesthesia during fUSi in head-fixed animals^75,76^ and recording the entire brain at high sampling rates thanks to the recent development of multi-array probes^77,78^.

To conclude, this study provides new insights into TSPO-dependent contributions to large-scale hippocampal network dysfunction in Alzheimer’s disease. Importantly, hippocampal abnormalities manifest at different timescales and locations along the ventral-dorsal axis, reflecting regional specializations in response to pathology. These results highlight fUSi as a powerful tool to bridge molecular pathology and systems-level dysfunction through integrated measurement of cerebral network activity at high spatiotemporal resolution and wide field-of-view, with the potential to guide mechanistic studies and therapeutic interventions. Furthermore, these findings indicate that modulation of TSPO and inflammatory pathways may offer a therapeutic strategy for AD by restoring brain activity and network integrity, independently of Aβ and tau pathology.

## Acknowledgements

V.Z. is supported by a Swiss National Science Foundation (SNSF) ECCELLENZA (PCEFP3_203005). S.T. is supported by a SNSF Ambizione Grant (223414).

## Statements and declarations

### Ethical considerations

Experimental approval was obtained from the ethics committee for animal experimentation of the canton of Geneva, Switzerland.

### Author contribution statement

AMB and BBT designed and performed the experiments. AMB and VD performed the preliminary analysis. BV and VZ analyzed the data. GS, ES, MK contributed to data acquisition. KC, LA, ST, SS, OB and PM contributed to data interpretation. BV, AMB, VZ and BBT wrote the manuscript. All authors reviewed and edited the manuscript.

### Disclosure of interest

The authors report no competing interests.

### Funding statement

This work was supported by the Swiss National Science Foundation (no. 310030-212322).

### Data availability

The fUSi data will be made publicly available upon final publication.

## Notes

### Competing Interest Statement

The authors have declared no competing interest.

