## Supplementary figures for "Minute- and second-scale hippocampal network dysfunctions in the 3xTgAD mouse model of Alzheimer’s disease are prevented by TSPO knockout"

### Supplementary information

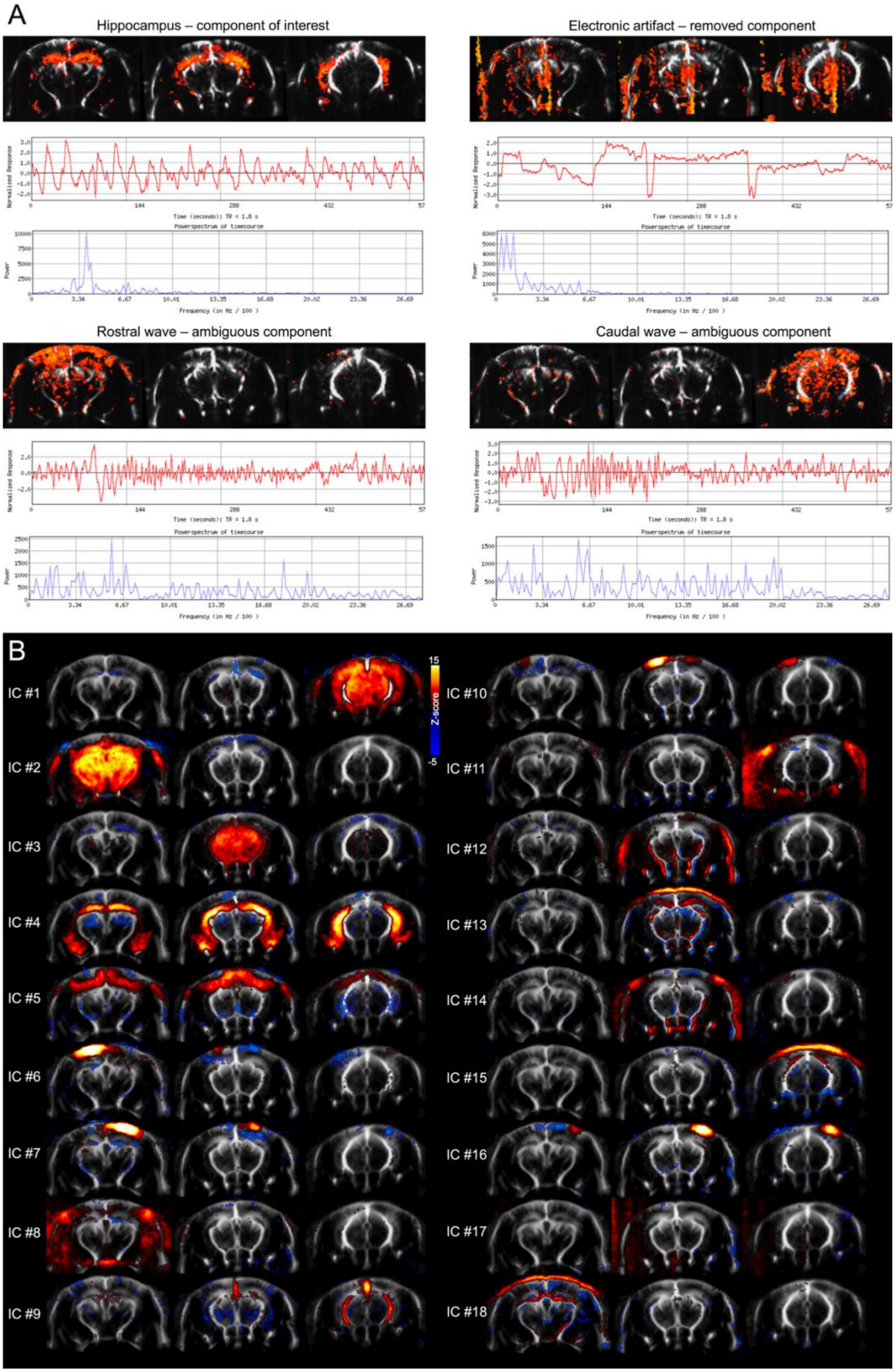
A
Hippocampus – component of interest
Electronic artifact – removed component
Rostral wave – ambiguous component
Caudal wave – ambiguous component
Normalised Response
Time (seconds); TR = 1.8 s
Powerspectrum of timecourse
Power
Frequency (in Hz / 100 )
B
IC #1
IC #2
IC #3
IC #4
IC #5
IC #6
IC #7
IC #8
IC #9
IC #10
IC #11
IC #12
IC #13
IC #14
IC #15
IC #16
IC #17
IC #18
15
Z-score
-5

**Figure S1: Individual component analysis results at the individual or group-level.** (A) Examples of individual maps following ICA on the same acquisition. The hippocampal network component is clearly detected at the individual level, spanning the dorsal and ventral areas and including parts of the amygdala, displaying slow oscillations around 0.04 Hz (top, left). A noisy component is also detected, characterized by vertical patterns appearing inside or outside of the brain, preferentially in zones with less signal, showing a time-course with sharp and unphysiological fluctuations (top, right). On bottom, ambiguous components were found, characterized by global waves occurring in most brain pixels of a particular coronal position and displaying irregular oscillations, varying in amplitude and frequency. Only the obvious noisy component on top-right was regressed out during the denoising step. (B) All of the 18 ICA maps obtained after group-level analysis of the entire dataset, including components of interest and other ambiguous, noisy or unilateral components.

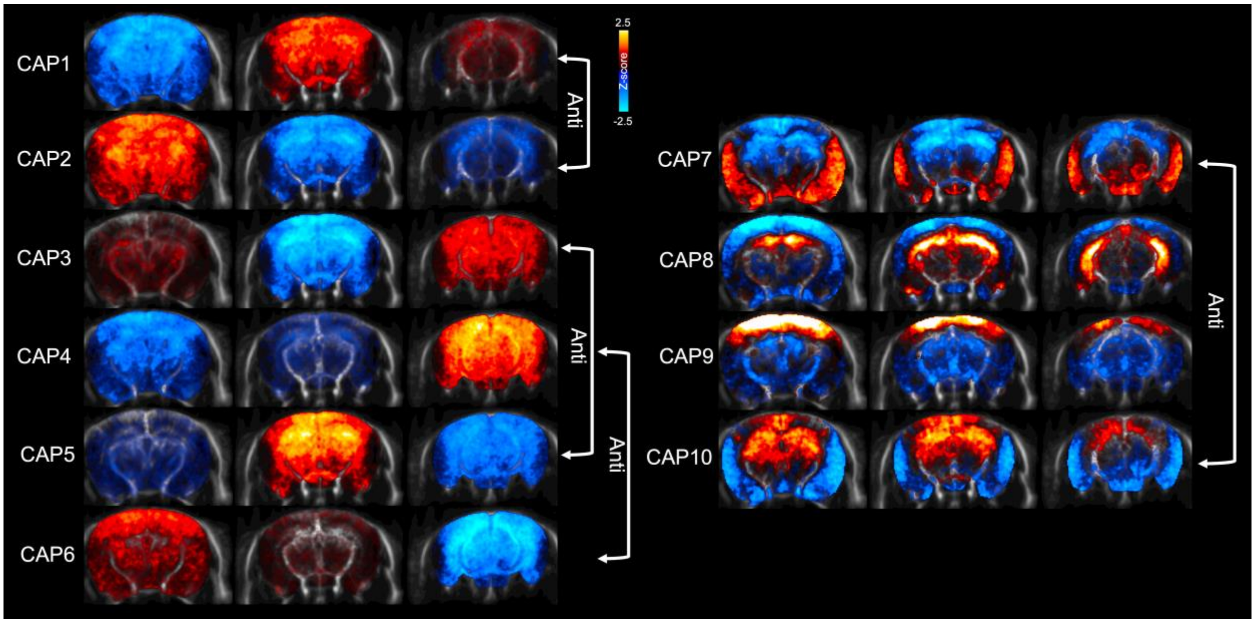


**Figure S2: Co-activation patterns group-level analysis.** All of the 10 CAPs maps obtained after group-level analysis of the entire dataset, including CAPs of interest on the right and global CAPs on the left, that typically involved signal increase in one coronal slice at a time with concurrent decrease in other slices. Some of the CAPs were almost exact opposites in terms of signal change polarity, such as CAP1 and 2, CAP3 and 5, CAP 4 and 6, or CAP 7 and 10.
